# Weak selection and the separation of eco-evo time scales using perturbation analysis

**DOI:** 10.1101/2021.09.14.460209

**Authors:** Philip Gerlee

**Affiliations:** Chalmers University of Technology

**Keywords:** eco-evo dynamics, pertubation analysis, time scales

## Abstract

We show that under the assumption of weak frequency-dependent selection a wide class of population dynamical models can be analysed using perturbation theory. The inner solution corresponds to the ecological dynamics, where to zeroth order, the genotype frequencies remain constant. The outer solution provides the evolutionary dynamics and corresponds, to zeroth order, to a generalisation of the replicator equation. We apply this method to a model of public goods dynamics and construct, using matched asymptotic expansions, a composite solution valid for all times. We also analyse a Lotka-Volterra model of predator competition and show that to zeroth order the fraction of wild-type predators follows a replicator equation with a constant selection coefficient given by the predator death rate. For both models we investigate how the error between approximate solutions and the solution to the full model depend on the order of the approximation, and show using numerical comparison, for ***k* = 1** and **2**, that the error scales according to ***ε***^***k*+1**^, where ***ε*** is the strength of selection and ***k*** is the order of the approximation.

## 1 Introduction

The processes that affect the size and composition of natural populations occur on a wide range of spatial and temporal scales. For example, speciation can occur due to spatial segregation of subpopulations, which genetically diverge due to adaptions to the local environment [1]. Spatial variation of the environment on smaller scales typically leads to trade-offs such as the growth-mortality trade-off among tropical trees which is caused by variation in light availability [2].

In terms of temporal effects it was previously held that measurable evolutionary change required vast amounts of time, on the order of thousands or millions of generations, to occur, and that therefore there was a clear separation between evolutionary and ecological dynamics. The latter occurs on the order of ten generations or less, but recent field studies and theoretical work has contradicted this clear cut separation and instead shown that evolution can occur over tens of generations, blurring the difference between evolutionary and ecological dynamics [3]. This rapid change can be driven by a changing environment, e.g. as in the industrial melanism in the peppered moth [4], or as in the case of Darwin’s finches by migration into a novel environment with many adaptive peaks [5].

The entanglement of eco-evo time scales raises the question of when it is valid to speak of evolutionary and ecological dynamics as two separate processes. One way to answer this question is to make use of perturbation analysis, which formalises the idea of processes acting on different time scales [6, 7], and makes it possible to quantify the error which the assumption of an absolute separation of time scales introduces. Perturbation analysis is applicable when a model contains a very small (or very large) parameter and aims at expressing the solution as a power series in this small parameter. The first term in this infinite series corresponds to an absolute separation of time scales, and higher order terms provide a correction to the zeroth order approximation. In ecology this method has been used in order to show that the logistic equation can be derived by assuming a separation of time scales between trophic levels, where the resource dynamics occurs on a much faster time scale compared to the population dynamics [8, 9]. The technique has also been used in evolutionary game theory [10], and more generally in stochastic models of evolution [11], in order to obtain analytical results for e.g. fixation probabilities.

Here we apply perturbation analysis to a general population dynamical model where the small parameter corresponds to the difference in frequency-dependent selection between different genotypes. We show that if selection is weak the model can be split up into distinct parts describing the evolutionary and ecological dynamics. We also apply the method to one model of public goods dynamics and one model of predator competition, and investigate how the error scales with the order of the approximation. Lastly, we discuss the relation between our framework and other methods for deriving approximate models in population dynamics.

## 2 General model

We consider *N* interacting subpopulations that represent distinct genotypes (or phenotypes). The population dynamics are assumed to be deterministic and driven exclusively by birth and death and we currently disregard the effects of migration and mutation, although these processes could be added to the model. The dynamics are then governed by the following system of coupled ordinary differential equations:

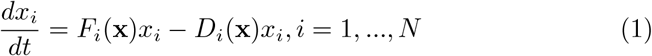

where *F*_*i*_(**x**) is the per capita birth rate and *D*_*i*_(**x**) is the per capita death rate of genotype *i*, which in this general setting depend on size of all subpopulations population **x** = (*x*_1_, *x*_2_, …, *x*_*N*_) We now assume that both the birth rate and death rate can be decomposed into the product of a frequency-dependent and a density-dependent factor, i.e.

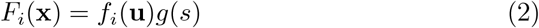

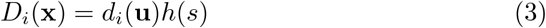

where the frequency-dependent factors *f*_*i*_ and *d*_*i*_ depend on the frequency of all types **u** = (*u*_1_, …, *u*_*N*_), where 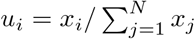 and the density-dependent factor depends on the total population size 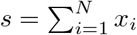, and are equal for all genotypes. We thus obtain the following system

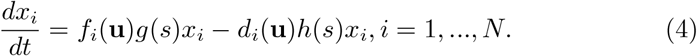

### 2.1 Change of variables

In order to get a better understanding of the dynamics of this system we make a change of variables to obtain a system of ODEs for the variables (**u**, *s*). This is related to the standard translation between the Lokta-Volterra equations and replicator dynamics (see Theorem 7.5.1 in [12]). Denoting time derivative with a dot, the rate of change of *u*_*i*_ can be written

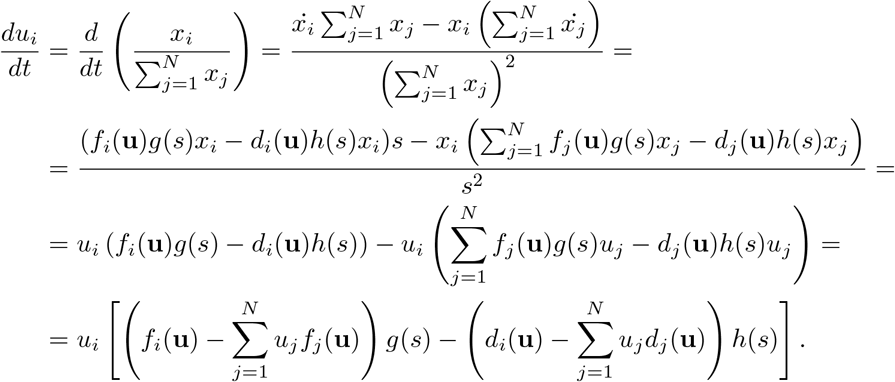

In order to simplify this expression we note that sums in the last expression are equal to the average frequency-dependent birth and death rates, which we denote:

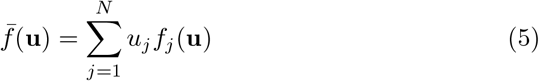

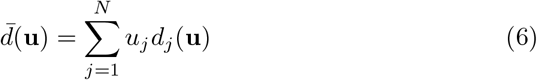

With these definitions we obtain:

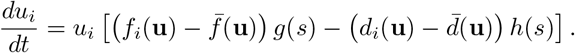

We now assume that the frequency-dependent birth and death rate for genotype *i* can be expressed as a baseline birth/death rate (common to all genotypes) plus a genotype-specific term of order *ε*:

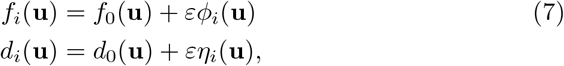

where *ε* ≪1 and *ϕ*_*i*_(**u**), *η*_*i*_(**u**) are some functions that depend on the frequency of all genotypes. The difference between the birth rate of genotype *i* and the mean birth rate can now be written: 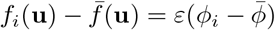, where 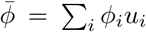. The assumption that *ε* ≪ 1 now implies that the difference in frequency-dependent birth rates between different genotypes is small, and since the same holds for the death rates selection is weak.

This results in the following equation for each *u*_*i*_:

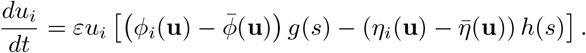

We now proceed to derive an equation for the total population size.

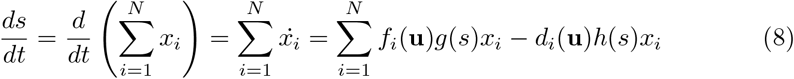

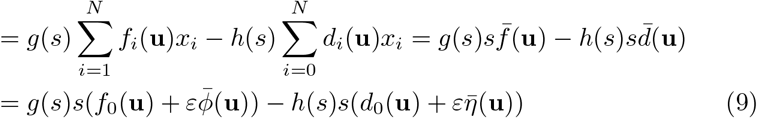

where in the second to last step we used the fact that *x*_*i*_ = *su*_*i*_. In summary we get the following coupled system of *N* + 1 equations:

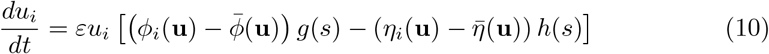

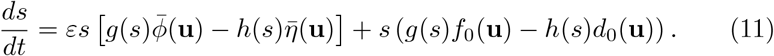

We thus have *N* + 1 equations that describe the dynamics in the (**u**, *s*)-space. However, note that one equation can be omitted since 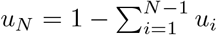.

### 2.2 Perturbation solution

We would now like to express the solutions *u*_*i*_(*t*) and *s*(*t*) as a power series in our small parameter *ε* (i.e. a perturbation solution):

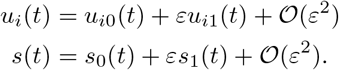

The next step is to insert this ansatz into (10), but here care has to be taken with expressions of the form *ϕ*_*i*_(**u**) and *g*(*s*), since we need to evaluate expressions of the form *ϕ*_*i*_(**u**_0_(*t*) + *ε***u**_1_(*t*) + 𝒪(*ε*^2^)). Now if the functions are multivariate polynomials it is readily seen that

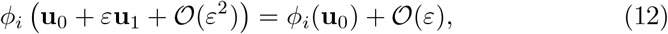

and similarly

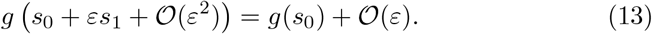

If our functions are not polynomials we can invoke the Stone-Weierstrass approximation theorem, which states that any continuous function defined on a compact subset *X* ⊂ ℝ^*N*^ can be uniformly approximated by polynomials to any degree of accuracy [13]. If this should be the case we simply replace our original functions with the approximating polynomials and then make use of the property (12) and (13).

We now insert the perturbation ansatz into (10) and comparing terms of order *ε*^0^ we get

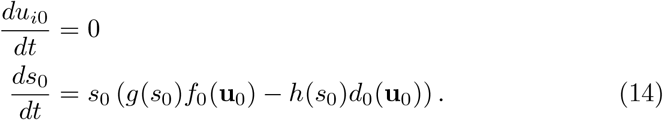

This implies that the frequency is unchanged and *u*_*i*0_(*t*) = *u*_*i*0_(*t* = 0) for all *t* ≥ 0, whereas the population size changes according to the difference between the density-dependent per capita growth rate *g*(*s*_0_) and death rate *h*(*s*_0_), each with a constant prefactor which depends on the baseline frequency-dependent birth/death rate of the initial frequency of genotypes **u**_0_.

This dynamics occurs on the 𝒪(1) time scale, and exclusively captures changes in population density. Therefore it is natural to define it as the ecological time scale. From the perspective of perturbation analysis this is usually referred to as the inner solution.

Higher-order corrections are obtained by comparing terms of order *ε* and higher, however this is not possible in this general setting where we have not specified the functions that describe frequency- and density-dependent birth and death rates. In the next sections, where we treat two specific models, higher-order terms are obtained.

In order to describe the dynamics for large times on the order of 1*/ε* (the outer solution) we define a new time scale *τ* = *εt* and express the solution as:

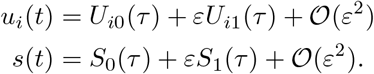

The time derivates on this time scale are given by

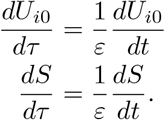

Using the power series ansatz yields, to order *ε*^0^

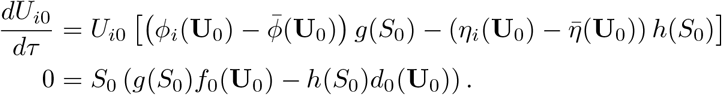

In order to eliminate *S*_0_ from the first equation we need to make two assumptions: firstly, assume that *S*_0_ ≠ 0 (i.e. the total population size is non-zero) and secondly assume that there exists an equilibrium density 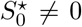 that solves the second equation *g*(*S*_0_)*f*_0_(**U**_0_) = *h*(*S*_0_)*d*_0_(**U**_0_). We also need to make sure that the equilibrium density is stable which can be determined by analysing the zeroth order equation for the density (14). The equilibrium is stable if the derivative of the right hand size evaluated at 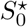 is negative:

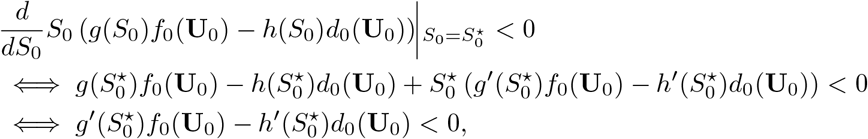

where the the last inequality is obtained since we assumed that 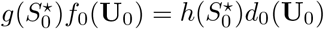. In summary the equilibrium density 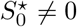 has to satisfy:

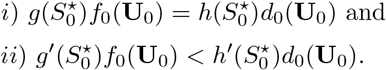

If multiple points exists that satisfy these conditions then the initial condition determines which equilibrium point is attained by the ecological dynamics.

If at least one such point exists we can write

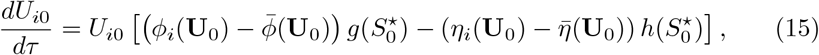

which describes the dynamics on the 1*/ε* time scale. On this time scale the population size equilibrates instantaneously and we only capture the changes in frequency. It is therefore natural to think of this as the evolutionary time scale.

In summary the perturbation analysis shows that the dynamics of the system (10) has two distinct regimes. For short times *t* = 𝒪(1) the population size *s* changes and the frequencies **u** are unchanged. This is typically referred to as the ecological time scale. For larger times *t* = 𝒪(1*/ε*) the density is constant whereas the frequency is altered. This is typically referred to as the evolutionary time scale.

## 3 Application to public goods game under logistic growth

We now apply the above results to a specific model, one that describes the dynamics of public goods production in a population consisting of producers and free-riders [14]. The model assumes that the public good is shared among all individuals and that it has a linear effect on the growth rate. This gives rise to the following frequency-dependent growth rates, where type 1 are the producers and type 2 are the free-riders (see [14] for details):

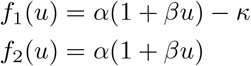

where *u* is the frequency of producers, *α* is growth rate in the absence of the public good, *β* describes impact of the public good and *κ* is the cost of production. In terms of assumption (7) we note that *f*_0_(*u*) = *α*(1+*βu*), whereas *ϕ*_1_ =− 1 and *ϕ*_2_ = 0, and as long as the cost of production is small (*κ* ≪ 1) selection is weak.

The two types are assumed to identical except for the public good production and therefore the the density-dependence is identical for both types. It is assumed to follow logistic growth with carrying capacity *K*, i.e. *g*(*s*) = 1 −*s/K*, and the per capita death rate is assumed to be constant and equal to *µ*. This implies that the dynamics of the population are given by:

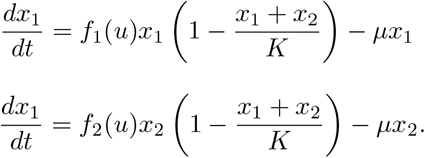

As before we change variables to *s* = *x*_1_ + *x*_2_ and 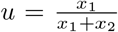 and obtain the following system

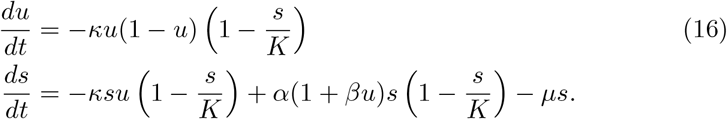

We now express the solutions *u*(*t*) and *s*(*t*) as a power series in *κ*:

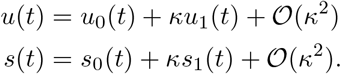

We insert the power series ansatz into (16) and gather terms with the same power of *κ*. To zeroth order in *κ* this yields

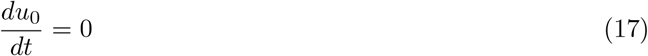

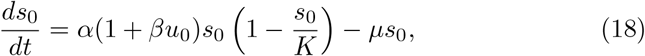

i.e. on the ecological time scale we obtain logistic growth with a growth rate that depends on the initial frequency of producers *u*_0_. With some additional algebra we obtain the following equations to first order in *κ*:

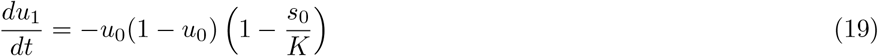

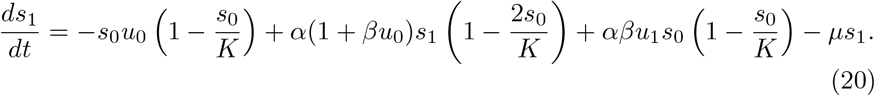

Here things get more complicated. Both the population size and frequency of producers change and the time derivative of *u* depends *s* and vice versa. Thus, we no longer have the separation of ecological and evolutionary time scales observed at zeroth order.

By carrying out the change of variables *τ* = *κt* we obtain the system:

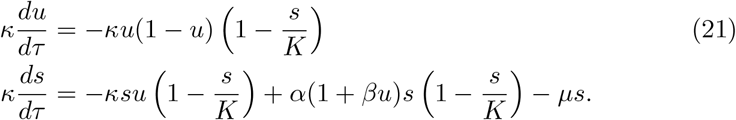

We now employ the outer power series ansatz

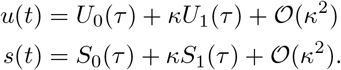

and obtain to zeroth order in *κ*:

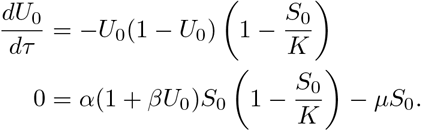

The second equation can be solved for *S*_0_ and ignoring the trivial solution *S*_0_ = 0 yields

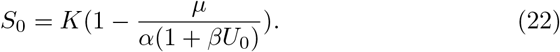

Inserting this into the first equation now gives

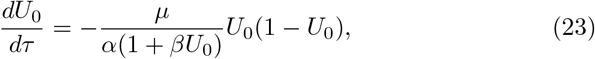

which provides us with the zeroth order dynamics on the evolutionary (1*/κ*) time scale. Comparing terms of order *κ* we obtain after some calculations:

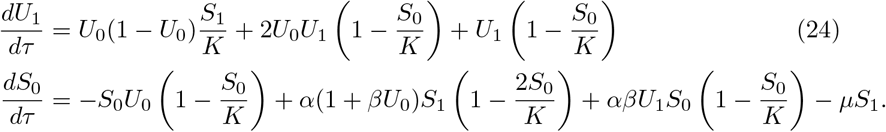

Since we know *S*_0_ from (22) we can solve the second equation for *S*_1_ and obtain

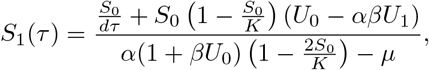

where

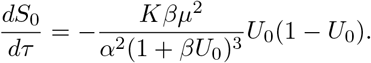

This expression for *S*_1_ can then be substituted into (24) to yield a closed ODE for *U*_1_.

### 3.1 Numerical comparison

In order to compare the accuracy of the zero order approximation of the original system (16) we calculate a composite solution using the method of matched asymptotic expansions [15]. The composite solution is obtained by adding the inner and outer solutions and subtracting the overlap, which is given by the the outer limit of the inner solution, and the inner limit of the outer solution. If we assume that the initial conditions are given by (*u*(*t* = 0), *s*(*t* = 0)) = (*a, b*), and since *u* remains unchanged for the inner solution, this implies that

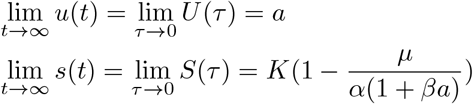

where the equality for *s* comes from the fact that *s* is completely determined in terms of *u* in the inner solution. The composite solution can now be written

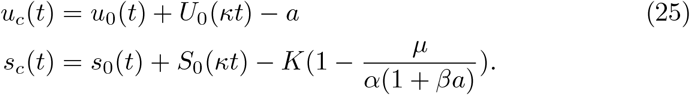

A numerical solution to (16) is shown in Figure 1A for different values of the small parameter *κ* together with the zero order composite solution (25). As expected a small value of *κ* leads to solution which shows clearer separation in ecological and evolutionary dynamics where the population first grows to carrying capacity without changes in frequency and then a subsequent change in frequency.

**Fig. 1.**
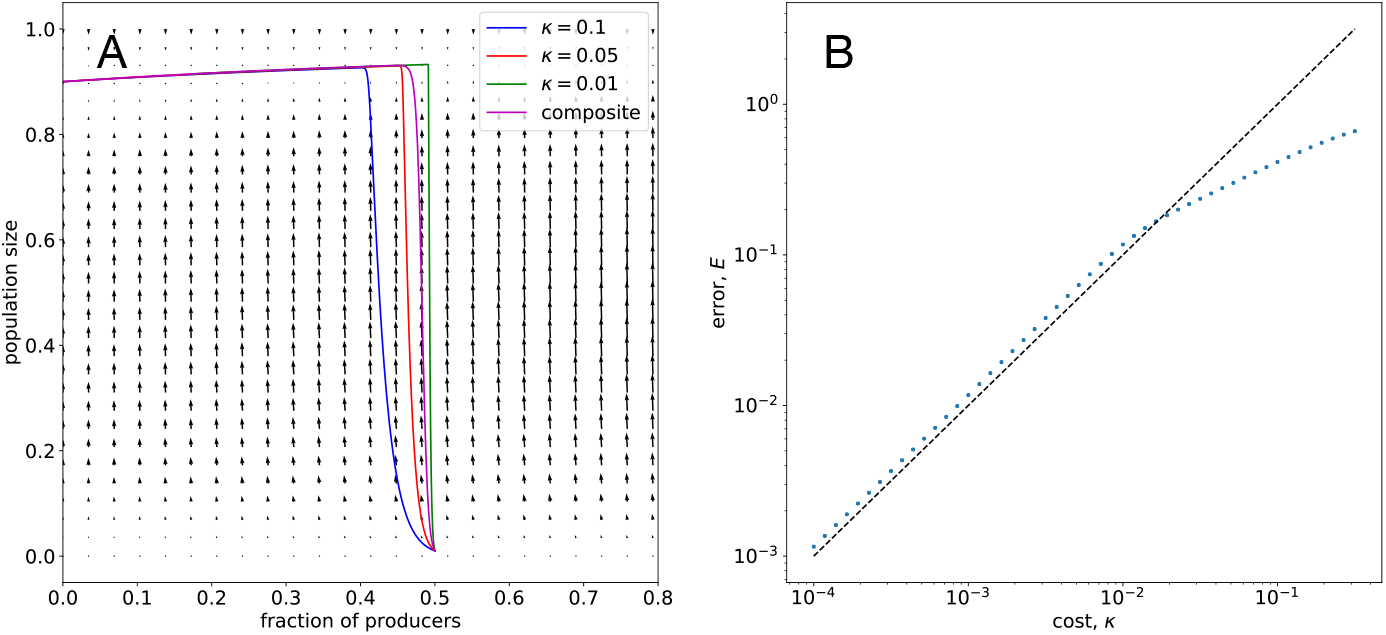
A Solution of (16) for different values of *κ* with the initial condition (*u, s*) = (0.5, 0.01). The other parameters are set to *α* = *β* = 1 and *K* = 1. The magenta curve shows the zeroth order composite solution (25). The underlying vector field corresponds to *κ* = 0.1. B The total error between the numerical solution of the original system (16) and the composite solution (25) as defined by (26) with *T*_*max*_ = 2000. The initial condition was set to (*u, s*) = (0.5, 0.01) and the parameters are set to *α* = *β* = 1 and *K* = 1. The dashed line corresponds to *E*(*κ*) ~ *κ*

In order to quantify the error introduced by the zero order approximation we calculate the total error

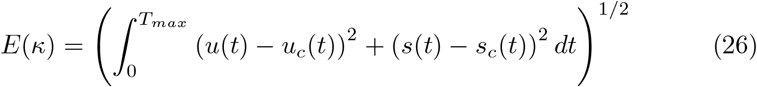

where all solutions are calculated numerically using the `lsoda`-solver as implemented in SciPy’s `odeint`. Figure 1B shows that as expected the error *E*(*κ*) of the zero order composite solution scales as *κ* as long as *κ* ≪ 1.

We also compare the accuracy of the zeroth and first order approximation of the population size on the ecological time scale by solving (18) and (20) numerically and calculating the error compared to the numerical solution of the full system. The initial conditions for the first order ODEs are chosen such that the approximation is valid for all values of *κ*, which corresponds to setting *u*_1_(*t* = 0) = *s*_1_(*t* = 0) = 0. The solutions are shown in figure 2A and the total error in 2B. This shows that the error scales according to *κ* for the zeroth order solution and *κ*^2^ for the first order solution.

**Fig. 2.**
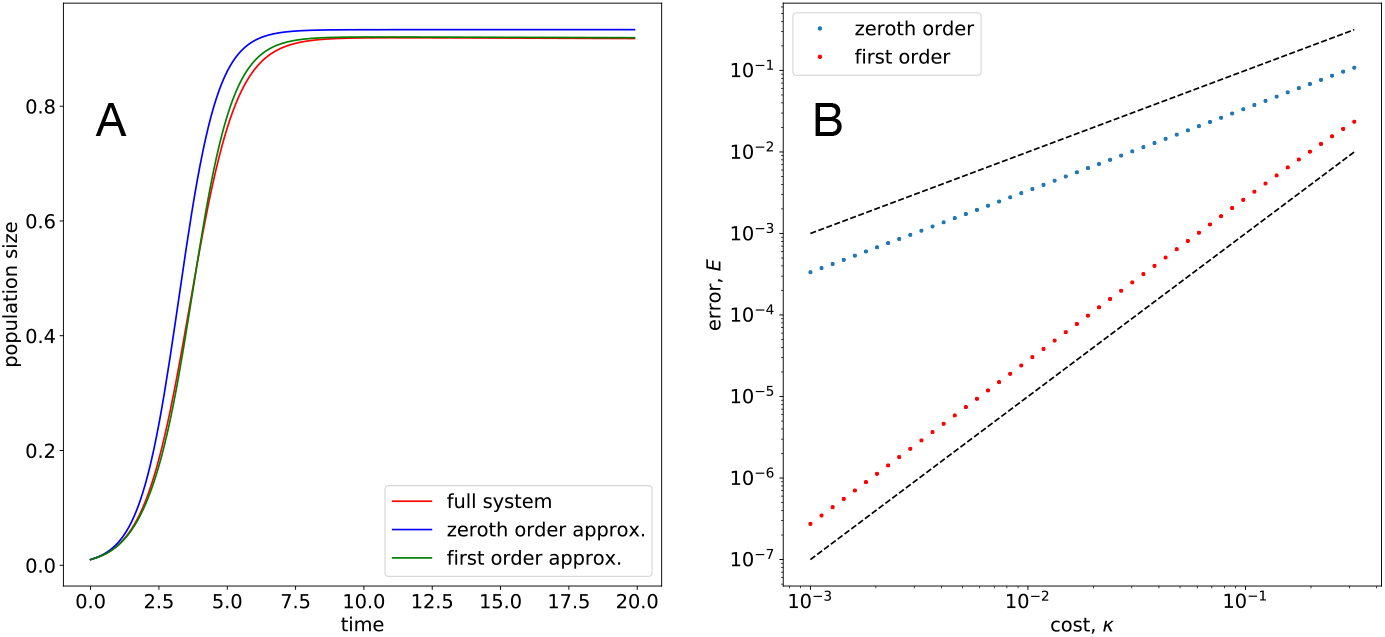
A Solutions of the ecological dynamics (18) and (20) for *κ* = 0.2 with initial condition (*u, s*) = (*u*_0_, *s*_0_) = (0.5, 0.01) and (*u*_1_, *s*_1_) = (0, 0). The other parameters are set to *α* = *β* = 1 and *K* = 1. B The total error for the zeroth and first order approximation. The dashed lines have slope 1 and 2 respectively.

## 4 Application to predator competition in a Lotka-Volterra system

We now turn to a more complicated system of a prey and two competing predators:

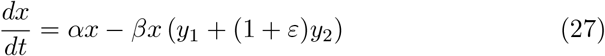

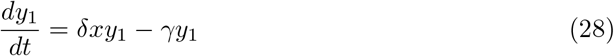

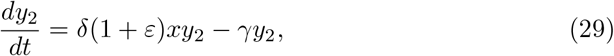

where *x* denotes the population size of the prey, *y*_1_ the population size of the wild-type predator and *y*_2_ the invading predator, which is a factor (1 + *ε*) better at feeding on the prey. The global stability of the steady states of this system has previously been investigated [16], and here we are concerned with the dynamics of the special case when *ε* ≪ 1.

Although the entire system does not conform to the general form of (4), it will be useful to carry out the outlined perturbation solution since it allows us to analyse a case where the ecological dynamics does not converge to a stable equilibrium point, but rather to a limit cycle.

We focus on the predator populations and therefore consider the dynamics of the total population of predators *s* = *y*_1_ + *y*_2_ and the fraction of the wildtype predator *u* = *y*_1_*/*(*y*_1_ + *y*_2_). In order to carry out the perturbation solution we make a change of variables and consider the system in terms of *x, s* and *u*. After some algebra we obtain:

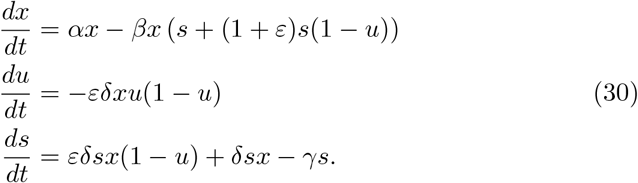

We begin with the perturbation solutions on the ecological time scale and make the ansatz:

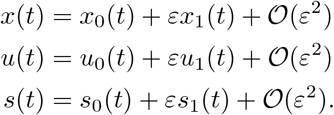

By inserting this into (30) and equating powers of *ε* we obtain to zeroth order

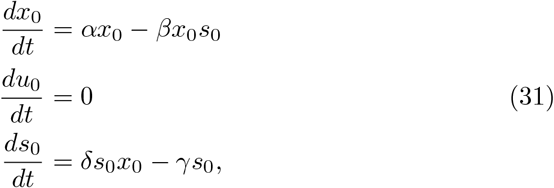

which we recognise as a standard Lotka-Volterra system in the variables (*x*_0_, *s*_0_), and where the fraction of wild-type predator *u*_0_ remains constant.

To first order in *ε* we obtain

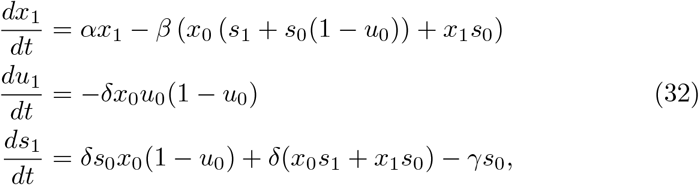

where we see a decrease in the fraction of the wild-type (since *x*_0_*u*_0_(1 −*u*_0_) *>* 0).

In order to analyse the evolutionary dynamics we carry out the change of variables *τ* = *εt* and obtain

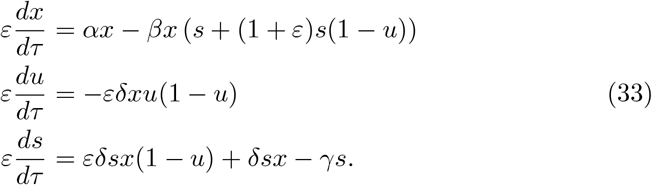

We then make the ansatz:

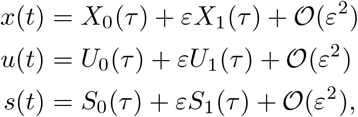

and obtain to zeroth order in *ε*

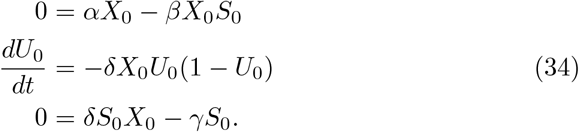

The first and third equation together yield *X*_0_ = *γ/δ* and *S*_0_ = *α/β*, which we recognise as the steady state of the zeroth order ecological dynamics (31). Thus, on an evolutionary time scale (to zeroth order) the prey population and the total predator population take on their ecological steady state values. The fraction of wild-type predators, on the other hand, follow a replicator-like equation, with a constant selection coefficient given by −*δX*_0_ = −*γ*, which is the death rate of the predators.

Comparing terms of order *ε* we obtain

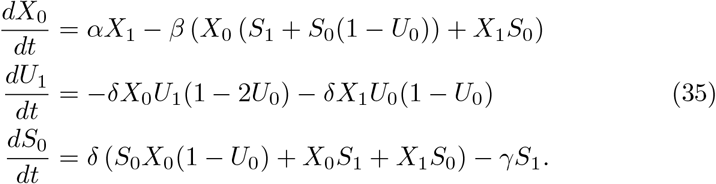

Since both *X*_0_ and *S*_0_ are constant, the first and third equation have a left hand side equal to zero, and can both be solved for *X*_1_ and *S*_1_ respectively. After some calculations we obtain 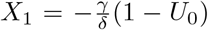 and 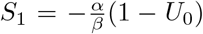. To first order on the evolutionary time scale we thus see a change in the prey and total predator population size according to

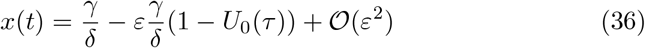

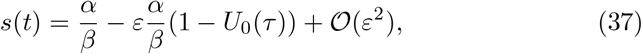

where the fraction of wild-type predators *U*_0_(*t*) changes according to (34) and the first order correction according to (35).

### 4.1 Numerical comparison

In this example where the ecological dynamics do not converge to a stable state (as it did for the public goods game), but instead enter into a limit cycle, we are not able to form a composite solution valid for all times. Instead we consider the dynamics on the ecological and evolutionary time scales separately. We focus on the fraction of wild-type predators and investigate how the zeroth and first order approximations on the ecological ((31) and (32)) and evolutionary ((34) and (35)) time scales compare to the solution of the full Lotka-Volterra system (30). All solutions were calculated numerically using the `lsoda`-solver as implemented in SciPy’s `odeint`. The result is shown in figure 3A where the inset shows the ecological dynamics (up to *t* ≈ 20) and the evolutionary dynamics is shown on the large axes. The full solution exhibits oscillatory dynamics and at the same decreases in a sigmoid fashion. The amplitude of these oscillations are diminished as *ε* decreases (data not shown). To zeroth order on the ecological time scale we obtain a constant fraction of wild-type predators, whereas the first order approximation tracks the oscillation fairly well approximately halfway through one cycle. On the evolutionary time scale the zeroth and first order approximations both capture the sigmoid decrease of the wild-type predator.

**Fig. 3.**
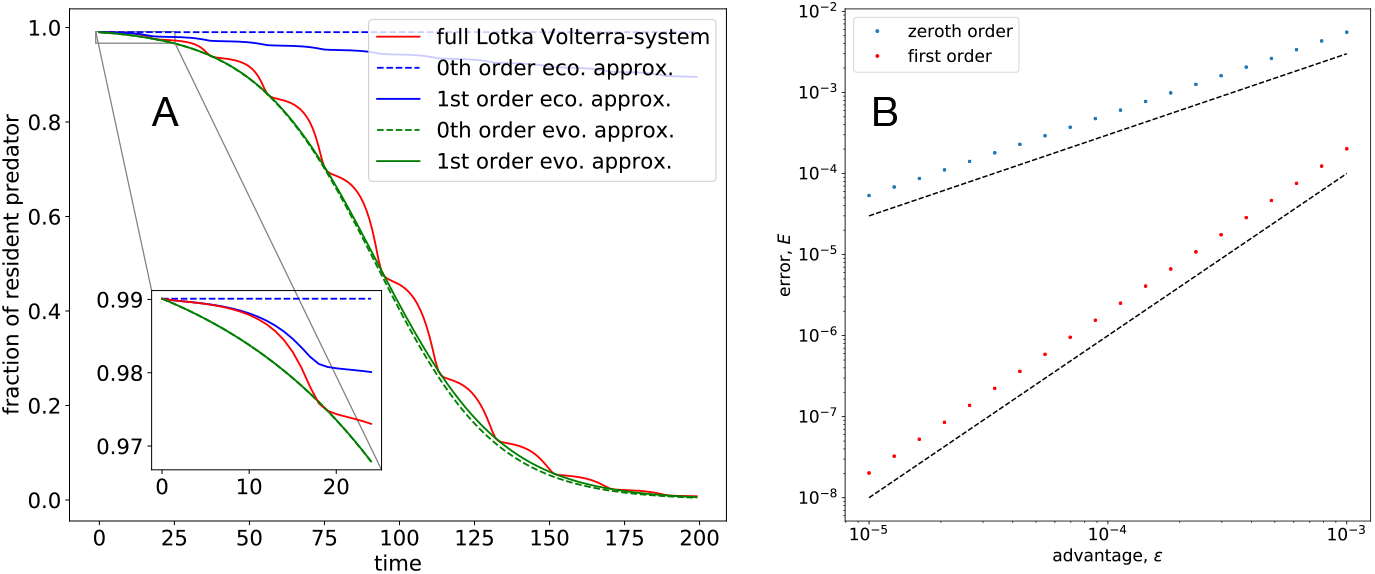
A The fraction of wild-type predators as a function of time for the full Lotka-Volterra system (red), on the ecological time scale (blue) and evolutionary time scale (green). Zeroth order approximations are dashed whereas first order approximations are shown as solid lines. The parameters are *ε* = 0.05, *α* = 0.2, *β* = 1, *δ* = 0.5 and *γ* = 1. The initial conditions are (*x, y*_1_, *y*_2_) = (1, 1, 0.01). B The total error between the numerical solution of the original system (30) and the zeroth and first order approximations on the ecological time scale (see inset in panel A). The total time was set to *T*_*max*_ = 20 and the parameters and initial conditions were set as in panel A. The dashed lines corresponds to *E*(*κ*) ~ *ε* and *ε*^2^.

When comparing the error between the full Lotka-Volterra system and the approximations on the ecological time scale we see as expected that the error of the zeroth order solution scales as *ε*, whereas the error of the first order solution scales as *ε*^2^.

## 5 Discussion

We have shown how perturbation analysis together with the assumption of weak selection provides a formal way of expressing the dynamics of a population on ecological and evolutionary time scales. The approximate solution is expressed as a power series in a parameter that quantifies the deviation between the frequency-dependent birth and death rates of each genotype and the average. We have applied the method to a specific model of a two strategy public goods game and the competition between two predators in a Lotka-Volterra model and showed numerically that the error introduced scales with the selection coefficient to the power *k* + 1, where *k* is the order of the approximation.

An important result is the derivation of the zeroth order equation for the outer/evolutionary dynamics (15), which can be viewed as a generalisation of the replicator equation

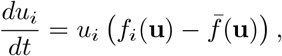

with the addition of density-dependent rates that modulate the intensity of selection. Our equation also captures the effects of frequency-dependence on both birth and death rates. The replicator equation has been derived for constant [17] and exponentially growing [18] populations, whereas (15) holds (to zeroth order in *ε*) for any type of density- and frequency-dependent growth.

The model of the public goods game has been analysed previously in order to obtain an approximate equation of the change in producer frequency [14]. In that case an equation for the invariant manifold was derived and solved in terms of a power series in the death rate *µ*. The resulting differential equation for the change in producer frequency was given by

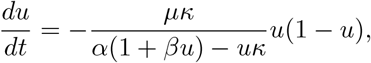

which at a first glance differs from (23). However, a Taylor expansion of the prefactor to first order in *κ* is given by 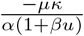 and if time is rescaled according to *τ* = *κt* (as was done to obtain (23)) we see that an equation identical to (23) is obtained. The two approaches thus yield identical equations for the evolution of the producer frequency.

We have also shown that our framework can be applied to models where the ecological dynamics follow oscillatory dynamics such as in the Lotka-Volterra system (30). Also here we obtain to zeroth order a replicator equation for the fraction of the wild-type prey (34), where the selection coefficient is constant and given by the death rate of the predators. We showed that higher order corrections to this simple relationship can be obtained, but also that they add little to the zeroth order solution, at least for small selection strengths. The Lotka-Volterra system is overly simplistic and does not account for mechanisms such as bounded growth of prey and predator handling-time. However, our framework is general and can also be applied to predator-prey models that are more realistic and thus more complex.

The concept of weak selection and the separation of ecological and evolutionary time scales which follows from it has been used extensively in mathematical modelling of population biology [19]. It is a central concept in adaptive dynamics, which is a framework that describes the long-term evolutionary dynamics of an evolving population [20]. There it is assumed that evolution follows a ‘trait substitution sequence’ in which monomorphic populations are invaded by fitter mutants that displace the wild-type. This process is captured by the so called canonical equation [21], which describes how the value of the traits under selection evolves over time, and is obtained as deterministic approximation of a directed random walk in the trait space. Notably this framework has been extended to situations where the ecological dynamics are oscillatory and chaotic [22].

Weak selection also features heavily in evolutionary game theory (EGT) since it simplifies the analysis and allows for explicit calculations of e.g. fixation probabilities [23]. The impact of higher order terms in EGT has been investigated by Wu et al. [10], who showed that the universality of weak selection breaks down when higher order terms are considered and that the microscopic model of evolution influences the dynamics. In addition is also also been shown that the ranking of strategies depends on the strength of selection and therefore that zeroth order approximations are a poor proxy under strong selection, in particular in multi-player games [24].

Related to the work on EGT is Allen and McAvoy’s analysis of fixation probabilities in populations of arbitrary size and spatial structure [11], where weak selection allows for computation of fixation probabilities in polynomial rather than exponential time. They have also considered a wide array of genetic structures (e.g. haploid, diploid etc.), and also there perturbation analysis as applied to selection strength plays a central role in obtaining analytical results [25].

The above mentioned EGT-models as well as those of Allen and McAvoy describe populations of a fixed size. This condition is relaxed in Aragasinski & Broom [26] who consider populations of varying size where birth and death rates are affected by both density- and frequency-dependence. They consider a system of coupled ODEs, which is a similar setting to ours, but are restricted to logistic density-dependence. Their treatment of time-scale separation is also far from rigorous and they only consider the zeroth order equilibrium solution to the ecological dynamics when expressing the evolutionary dynamics. Variable population sizes are also considered in Cressman & Garay [27], which investigate stability in *N* − species coevolutionary systems. They claim that time scale separation is not required in their analysis, and instead their focus is on ‘stationary density surfaces’ which correspond to equilibria of the ecological dynamics. However, such equilibria are not reached (for general initial conditions) unless the ecological time scale is completely separate from the evolutionary. Another model that considers a variable population size is analysed by Parsons & Quince [28], which is formulated in terms of a two-dimensional Markov chain, where the transition rates contain a logistic density-dependence. They also make use of perturbation analysis in order to calculate fixations probabilities, but in their analysis it is the reciprocal of the carrying capacity which plays the role of the small parameter.

Lastly, it should be mentioned that weak selection has also been used in inclusive fitness theory, where it simplifies the calculations of fixation probabilities and selection gradients [29], and in the theory of branching processes [30].

In contrast to the above mentioned contributions to our understanding of weak selection and its implications for eco-evo dynamics we would like to highlight that the work presented here is the first to jointly consider: (i) general density-dependent growth (as opposed to e.g.[26]), (ii) a rigorous treatment of ecological and evolutionary time scales in terms of inner and outer solutions including composite solutions obtained using matched asymptotic expansions, and (iii) a quantification and numerical verification of the error introduced by using zeroth and first order approximations.

One limitation of our framework is that it only applies to deterministic models. For small populations stochastic models are more relevant since they account for demographic noise, and extending our methodology to such models would therefore be desirable. One way of achieving this could be to formulate the dynamics as an *N* − dimensional Markov chain that describes the population size of the considered genotypes and writing down the corresponding Fokker-Planck equation as in [28]. By assuming weak frequency-dependence and expanding the solution of the Fokker-Planck equation in a power-series of the selection strength it might be possible to obtain approximate solutions. Another, possibly simpler approach, would be to take the continuum limit of the Markov chain to obtain a system of *N* coupled stochastic differential equations [31]. In this setting questions about fixation probabilities and fixation times can be pursued using perturbation analysis coupled with the toolbox of stochastic analysis.

The framework we have introduced is applicable to a large class of population dynamical models with density- and frequency-dependence, and makes it possible to derive approximate solutions in the case of weak selection. The use of composite solutions in addition makes it possible to obtain solutions valid for all times. If selection is not weak it is possible to include higher-order terms in the power series expansion and still obtain distinct models for the ecological and evolutionary time scales. As was seen for the public goods and Lotka-Volterra model these higher order corrections typically involve changes in both density and frequency.

The method presented here also makes it possible to test if, for a given model, separation of time scales is a sensible approximation. This could for example be used to investigate if evolutionary game theory, which only describes changes in frequency, is a reasonable description of a certain system, or if changes in density have to be accounted for.

